# Dissecting the regulatory roles of ORM proteins in the sphingolipid pathway of plants

**DOI:** 10.1101/2020.08.24.264705

**Authors:** Adil Alsiyabi, Ariadna Gonzalez Solis, Edgar B Cahoon, Rajib Saha

## Abstract

Sphingolipids are a vital component of plant cellular endomembranes and carry out multiple functional and regulatory roles. Different sphingolipid species confer rigidity to the membrane structure, facilitate trafficking of secretory proteins, and initiate programmed cell death. Although the regulation of the sphingolipid pathway is yet to be uncovered, increasing evidence has pointed to orosomucoid proteins (ORMs) playing a major regulatory role and potentially interacting with a number of components in the pathway, including both enzymes and sphingolipids. However, experimental exploration of new regulatory interactions is time consuming and often infeasible. In this work, a computational approach was taken to address this challenge. A metabolic network of the sphingolipid pathway in plants was reconstructed. The steady-state rates of reactions in the network were then determined through measurements of growth and cellular composition of the different sphingolipids in Arabidopsis seedlings. The Ensemble modeling framework was modified to accurately account for activation mechanisms and subsequently used to generate sets of kinetic parameters that converge to the measured steady-state fluxes in a thermodynamically consistent manner. In addition, the framework was appended with an additional module to automate screening the parameters and to output models consistent with previously reported network responses to different perturbations. By analyzing the network’s response in the presence of different combinations of regulatory mechanisms, the model captured the experimentally observed repressive effect of ORMs on SPT. Furthermore, predictions point to a second regulatory role of ORM proteins, namely as an activator of class II (or LOH1 and LOH3) ceramide synthases. This activating role was found to be modulated by the concentration of free ceramides, where an accumulation of these sphingolipid species dampened the activating effect of ORMs on ceramide synthase. The predictions pave the way for future guided experiments and have implications in engineering crops with higher biotic stress tolerance.

**Author summary:** Due to their vital functional and regulatory roles in plant cells, increasing interest has gone into obtaining a complete understanding of the regulatory behavior of the sphingolipid pathway. However, the process of identifying new regulatory interactions is time consuming and often infeasible. To address this issue, ensemble modeling was used as an *in silico* method to test the ability of different regulatory schemes to predict all known pathway responses in a thermodynamically consistent manner. The analysis resulted in a significant reduction in the number of possible regulatory interactions. Mainly, the model predicts regulatory interactions between ceramides, ORMs, and ceramide synthases (especially class II). This framework can pave the way for biochemists to systematically identify plausible regulatory networks in understudied metabolic networks where knowledge on the underlying regulatory mechanisms is often missing. As future experimental works explore these predictions, an iterative cycle can begin wherein model predictions allow for targeted experiments which in turn generate results that can be reincorporated into the model to further increase prediction accuracy. Such a model-driven approach will significantly reduce the solution space traversed by the experimentalist.

## Introduction

Sphingolipids are a diverse group of membrane lipids essential in eukaryotic organisms. In plants, sphingolipids comprise up to 40% of the plasma membrane and are abundant components of cellular endomembranes such as the endoplasmic reticulum (ER), Golgi, and tonoplast [1,2]. Through their unique structural features, sphingolipids carry out several essential functions in plant cells. Glycosylated sphingolipids like glucosylceramides (GlcCer) and glycosylinositolphosphoceramides (GIPCs) contribute to membrane function and are involved in the trafficking of secretory proteins out of the cell [3,4]. The accumulation of other sphingolipids, namely long chain bases (LCBs) and ceramides, signal the initiation of programmed cell death (PCD) when plant cells are under environmental stresses, such as the presence of bacterial or fungal pathogens [5,6].

The sphingolipid biosynthesis pathway starts in the ER with the condensation of palmitoyl-CoA and serine to produce 3-ketosphinganine. This reaction is the first committed step towards sphingolipid LCB biosynthesis and is also the rate-limiting step of the sphinolipid biosynthetic pathway [7]. 3-Ketosphinganine is then reduced to sphinganine, which is the basic LCB. Sphinganine can then go through multiple modifications including unsaturation in Δ4 or Δ8 positions; hydroxylation at C4 and/or phosphorylation at its C1 position. LCBs can then be linked to a fatty acyl-CoA, typically with chain-lengths ranging from 16 to 26carbon atoms, to produce ceramides [8,9] by the activities of ceramide synthases. In Arabidopsis, two classes of ceramide synthases were identified. Class I, encoded by Longevity Assurance Gene One Homolog2 (*LOH2*), mostly operates on acyl-CoAs of length 16 and dihydroxy LCBs and class II, encoded by *LOH1* and *LOH3*, act on acyl-CoAs containing more than 22 carbons, also referred to as very long chain fatty acids (VLCFAs) and tri-hydroxy LCBs [9,10]. The ceramide backbone is further modified by glycosylation at its C-1 position to form GlcCer or linked to inositol phosphate and further glycosylated in Golgi bodies to yield GIPCs, the most abundant glycosphingolipid in plant cells [1,11,12].

Similar to many other biochemical pathways, the first committed step catalyzed by serine palmitoyltransferase (SPT) constitutes the main regulatory point in the pathway [7]. Tight regulation on SPT ensures sufficient production of sphingolipid components to maintain cellular growth, and simultaneously prevents the accumulation of PCD-inducing components under non-stress conditions. A small polypeptide of 56 amino acids referred to as the small subunit of SPT (ssSPT) is necessary for optimal activity of this enzyme [13]. Recently, orosomucoid-like proteins (or ORMs) have emerged as negative regulators of the sphingolipid pathway [14]. Studies conducted in *Arabidopsis thaliana* (hereafter Arabidopsis), *Saccharomyces cerevisiae* and mammalian cells have shown that the lack of functional ORM proteins results in accumulation of sphingolipids, especially ceramides and LCBs [14–17]. Interestingly, these proteins are essential to complete a life cycle in the model multicellular organisms Arabidopsis and mouse [16,17]. Although the exact regulatory mechanisms remain unknown, it has been shown that the ORM-SPT physical interaction is necessary to downregulate the activity of the enzyme [16,18].

Interestingly, in addition to the regulation at the first step of the biosynthetic pathway, ORMs were proposed to act as modulators of ceramide synthases [19]. It was shown that the overexpression of *ORM* genes leads to differential activity between the two classes of ceramide synthase. Activity of class I ceramide synthase (LOH2) was observed to decrease, while class II (LOH1/LOH3) activity was stimulated [19]. Conversely, downregulation of *ORM* gene expression produced the opposite effect. In addition, studies conducted on yeast and mammalian cells postulated a ceramide-ORM feedback regulation, in which the physical interaction between ceramides and ORMs leads to a decrease in SPT activity [20–22]. Taken together, these experimental observations started to uncover the global regulatory role played by ORMs in the sphingolipid pathway, however, there still remain critical knowledge gaps. For example, it is still not known whether ORMs have any regulatory interactions with the ceramide synthases and whether or not any other sphingolipid components interact with ORMs [19]. Furthermore, while several potential regulatory schemes were proposed to explain each of the observed pathway behavior, experimentally verifying each of the proposed mechanisms remains an impractical task.

Computational systems biology is a field concerned with modeling complex biological systems to test the feasibility of a predefined hypothesis or to generate new testable hypotheses based on existing observations [23]. A widely applied modeling framework that is used to address such challenges is kinetic modeling [24]. This framework describes the metabolic and regulatory processes occurring in the system through kinetic expressions (e.g. mass action, Michaelis-Menten). Starting with a set of initial conditions, the temporal behavior of the metabolic and regulatory network can be determined [25]. However, practical application of this framework is dependent on the availability of measured enzyme kinetic parameters for all enzymes in the network, which is usually not feasible. To overcome this limitation, ensemble kinetic modeling (EM) [26] was developed to sample through the entire allowable kinetic solution space to generate an ensemble of kinetic models that describe the system (see methods). This ensemble is then filtered using prior knowledge of the system’s response to different genetic perturbations [27]. EM was used to capture the inherent non-linearity of metabolic systems and to identify metabolic bottlenecks in the production of various industrially relevant compounds [28]. In addition, EM was used to predict the presence of regulatory interactions occurring in biochemical pathways [29]. Implementation of such procedure can therefore greatly accelerate the process of hypothesis generation and testing by predicting plausible regulatory mechanisms that satisfy the observed experimental behavior of the pathway under different growth conditions. This analysis can subsequently be followed up with experimental testing for validation and iterative model curation.

A number of studies have used different forms of kinetic modeling to study sphingolipid metabolism in yeast and mammalian cells [30–33]. These studies used different experimental data sets such as lipidomics, transcriptomics, and fluxomics to parametrize a predefined kinetic model describing the sphingolipid pathway. In most cases, the implemented methods assumed that the sphingolipid regulatory network was completely understood [30–32]. Furthermore, they did not account for the possibility that more than one parameter set can satisfy the observed experimental measurements. A recent study aimed at analyzing sphingolipid metabolism in yeast addressed these issues by implementing a new framework called inverse metabolic control analysis (IMCA) [33]. The authors used this framework to identify the key enzymes responsible for the observed sphingolipid profile. However, the study did not incorporate any regulatory proteins such as ORMs into their analysis.

In this work, a kinetic model of the Arabidopsis sphingolipid pathway was constructed to predict the pathway’s response to various internal and external perturbations. By using the EM approach, we identified a highly plausible regulatory interaction between ORM proteins and different components of the sphingolipid network. These interactions were deemed plausible as their presence was required in order to obtain the experimentally observed behavior of the system. By testing different sets of regulatory mechanisms, the implemented framework was used to eliminate postulated interactions which were not consistent with the experimentally observed response of the network to genetic perturbations. The novel prediction made by the analysis is the presence of a ceramide-ORM-ceramide synthase (class II) regulatory interaction. In this scheme, ORMs activate class II ceramide synthases (LOH1/LOH3); however, upon ceramides accumulation ORMs are repressed and not able to activate CS II causing a buildup of long chain bases (LCBs). These results illustrate how kinetic models can be used to predict regulatory mechanisms in biochemical systems despite lack of prior knowledge on enzyme kinetics. These predictions will result in a metabolic and regulatory model of the sphingolipid network capable of more accurately predicting the pathway’s response to different genetic manipulations as well its response to biotic and abiotic stresses.

## Results

### Metabolic network reconstruction

The reconstructed network comprises a knowledgebase of all the active metabolic transformations involved in sphingolipid biosynthesis in Arabidopsis (see S1 File). The network comprises 78 reactions associated with 24 genes. This resulted in an elementary reaction network consisting of 374 elementary reactions, excluding the elementary steps incorporated to describe regulatory mechanisms (see Materials and method). The large difference between the number of reactions and number of genes is due to the enzyme promiscuity present in the sphingolipid synthesis pathway. In Arabidopsis, the activity of the Δ4 desaturase is minimal in most tissue types [34], therefore this reaction was not included into the network. Moreover, the majority of acyl-CoAs found in sphingolipids are either C16 or C24 fatty acids [8,35]. Hence, the relatively negligible concentrations of other chain-length fatty acids were combined with one of these two groups based on their chain length. This was done to reduce redundancy in the added reactions and to avoid having an exaggerated number of kinetic parameters, which could potentially lead to overfitting.

Fig 1 shows the metabolic map used to construct the kinetic model. Several assumptions had to be made on the order of modifications occurring to the different components in the sphingolipid pathway. It was assumed that the enzyme LCB C4 hydroxylase (SBH1 and SBH2) mainly hydroxylates free long chain bases instead of ceramides containing dihydroxylated LCBs. This was based on previous evidence which showed that the *sbh1 sbh2* double mutant accumulated sphingolipids enriched in C16 fatty acids with dihydroxy LCBs coming from the ceramide synthase class I [36]. Moreover, based on the findings reported by Konig et al. [37], it was assumed that the acyl-hydroxylating enzyme fatty acid α-hydroxylase (FAH1 and FAH2) converted only ceramide-bound fatty acids and not free fatty acids. LCB Δ8 desaturase (SLD1 and SLD2) was also assumed to produce unsaturated LCBs bound to ceramides and not free LCBs [38,39]. Again, this was done to avoid redundancy, as omitting these assumptions would have led to an exponential increase in the number of reactions and therefore fitting parameters. In addition, it was assumed that sphingolipid turnover rates were negligible compared to their rate of formation. This assumption was necessary to calculate reference steady-state fluxes. The linear GIPC forming pathway involving IPC-synthases and several glycosyl or glucuronyl-transferases [40] was lumped into one step, as no information was present on the relative kinetics of the participating enzymes. Finally, since the majority of the glycosylated sphingolipids (GlcCers and GIPCs) contain ceramides with hydroxylated fatty acids, no reactions were added to convert non-hydroxylated ceramides into these complex forms.

**Fig 1.**
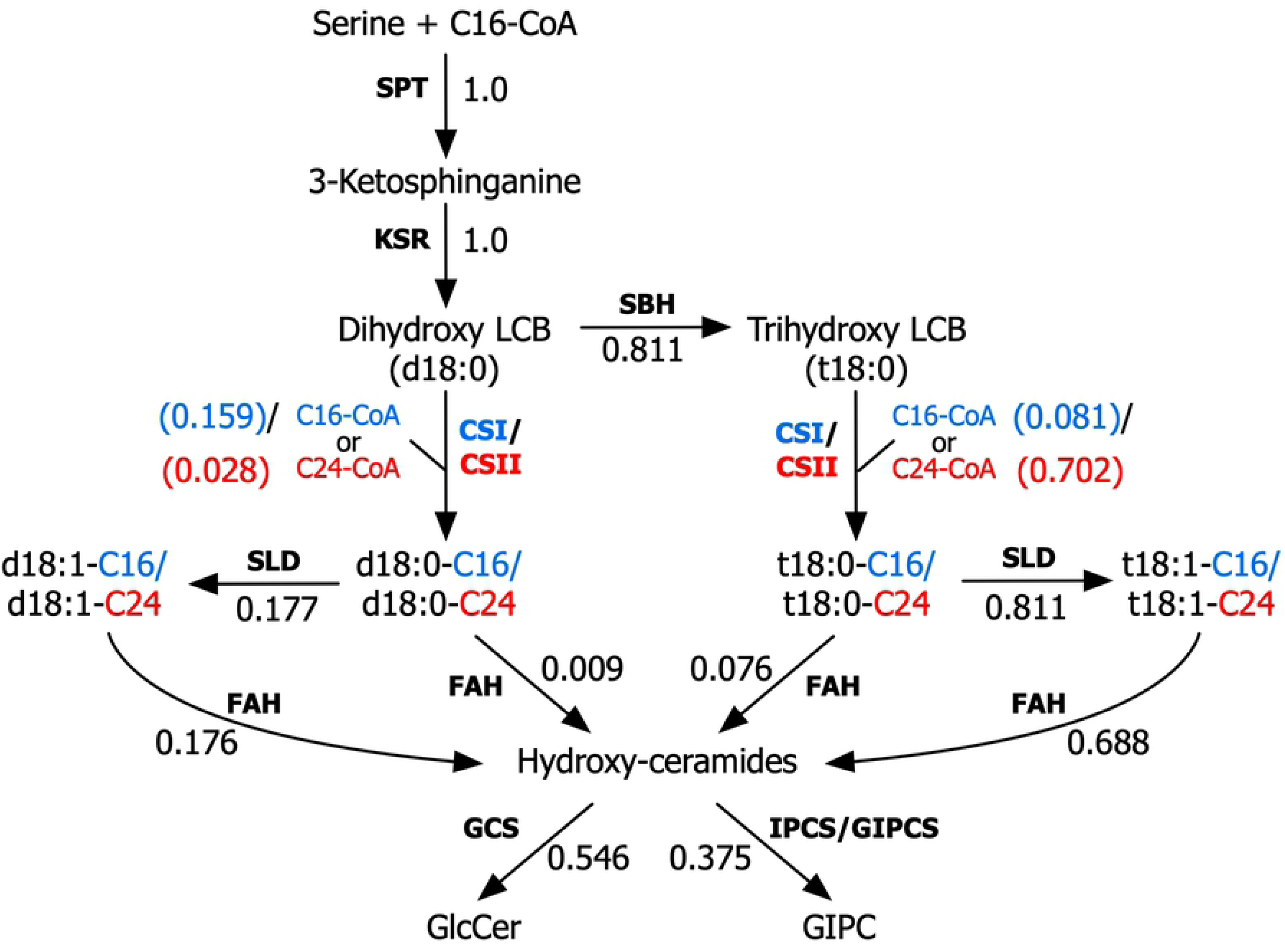
Simplified depiction of the sphingolipid pathway. A metabolic network displaying the main reactions constituting the sphingolipid pathway. Blue flux values correspond to reactions catalyzed by CSI. SPT: serine palmitoyltransferase, KSR: 3-ketosphinganine reductase, SBH: LCB C-4 hydroxylase, CSI: class I ceramide synthase, CSII: class II ceramide synthase, SLD: LCB Δ8 desaturase, FAH: fatty acid hydroxylase, GCS: glucosylceramide synthase, GlcCer: glucosylceramide, GIPCS: glycosyl inositolphosphoceramide synthase, LCB: long chain base. C16 and C24 are fatty acids of length 16 and 24 carbons, respectively.

### Generation of kinetic parameters

The Ensemble modeling framework requires the input of both the reference steady-state fluxes and the standard Gibbs free energies of the modeled reactions to generate the initial set of kinetic parameters [26] (see Fig 2). The fluxes were determined by measuring the growth rate and sphingolipid profiles of Arabidopsis seedlings at multiple time points during exponential growth (see materials and methods). By assuming negligible sphingolipid turnover and a constant sphingolipid composition, measurements of growth and composition were used to determine the accumulation rate of each sphingolipid component. Flux balance analysis (FBA) [41] was subsequently used to calculate the fluxes of the internal reactions in the network (see materials and methods). The values displayed in Fig 1 are normalized to SPT to illustrate how the flux going through this reaction is branched into the different sphingolipid products. As can be seen, the majority of the produced LCBs (73%) are channeled through class II ceramide synthase to produce VLCFA containing ceramides. Furthermore, under normal conditions, practically all of the sphingolipid components are non-phosphorylated (<1% phosphorylation). The activity of the complex sphingolipid producing enzymes GCS and GIPCS were similar. However, whereas the majority of GIPC products were VLCFA-containing ceramides, 30% of the flux through GCS went towards synthesizing C16-containing ceramides. This was an interesting observation for the wild type, since previous CSI knockout studies found that elimination of the C16-ceramide producing ceramide synthase has no observed change in phenotype under normal growth conditions [8].

**Fig 2.**
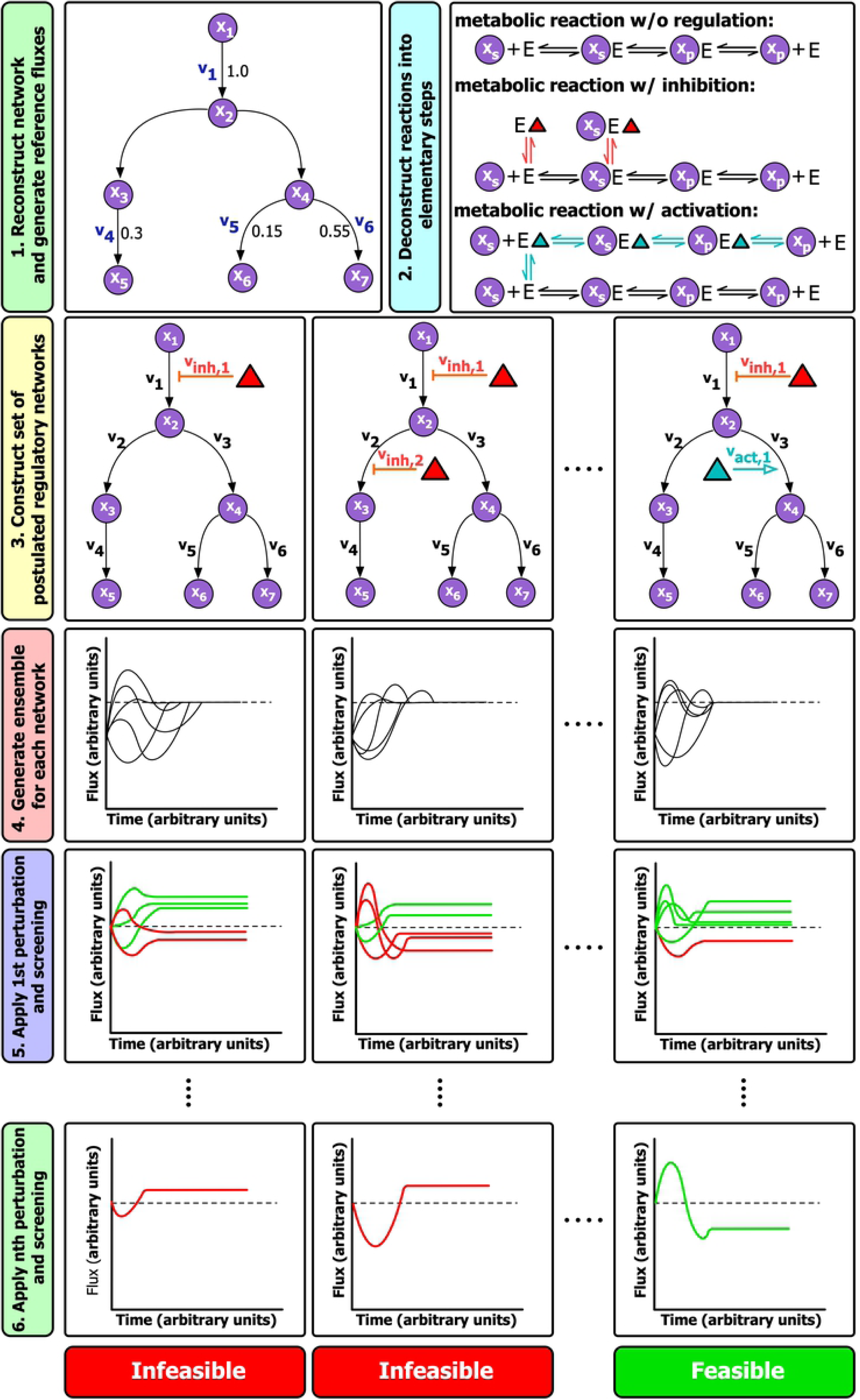
Workflow for using Ensemble modeling to predict regulatory mechanisms.

The component contribution method [42] was subsequently used to calculate the standard Gibbs free energy of each reaction in the pathway. As can be seen from the values in table 1, the standard free energy for most of the biochemical transformations in the pathway are negative and relatively far from equilibrium. Therefore, these reactions have a wide range of substrate/product concentration ratios that can result in an overall forward flux of the reaction [43] (see S2 File). However, the first and rate limiting step of the pathway, the reaction catalyzed by SPT [1], has a slightly positive standard free energy. This means that a positive substrate to product concentration ratio needs to be maintained in order for the reaction rate to stay in the forward direction. Furthermore, the rate of the reaction is directly affected by any changes in the actual free energy, and therefore in the concentrations of the participating reactants [43]. Once the required inputs were determined, a MATLAB implementation of the ensemble modeling framework was customized to generate the kinetic parameters (see materials and methods). This process was repeated for each of the tested regulatory schemes. Fig 3 illustrates how the generated set of kinetic parameters result in the measured steady-state fluxes for the reference (wild type) steady-state and how the generated models are screened based on their response to different perturbations.

**Table 1.** Standard Gibbs free energy values for reactions in the sphingolipid pathway.

| Reaction Name | $\Delta G^{\circ}$ (kJ/mol) |
| --- | --- |
| SPT | 8.9 |
| KSR | -17.1 |
| SBH | -397.4 |
| CS1 | -25.5 |
| CS2 | -27.6 |
| FAH | -93.0 |
| GCS | -4.5 |
| GIPCS | -5.1 |

**Fig 3.**
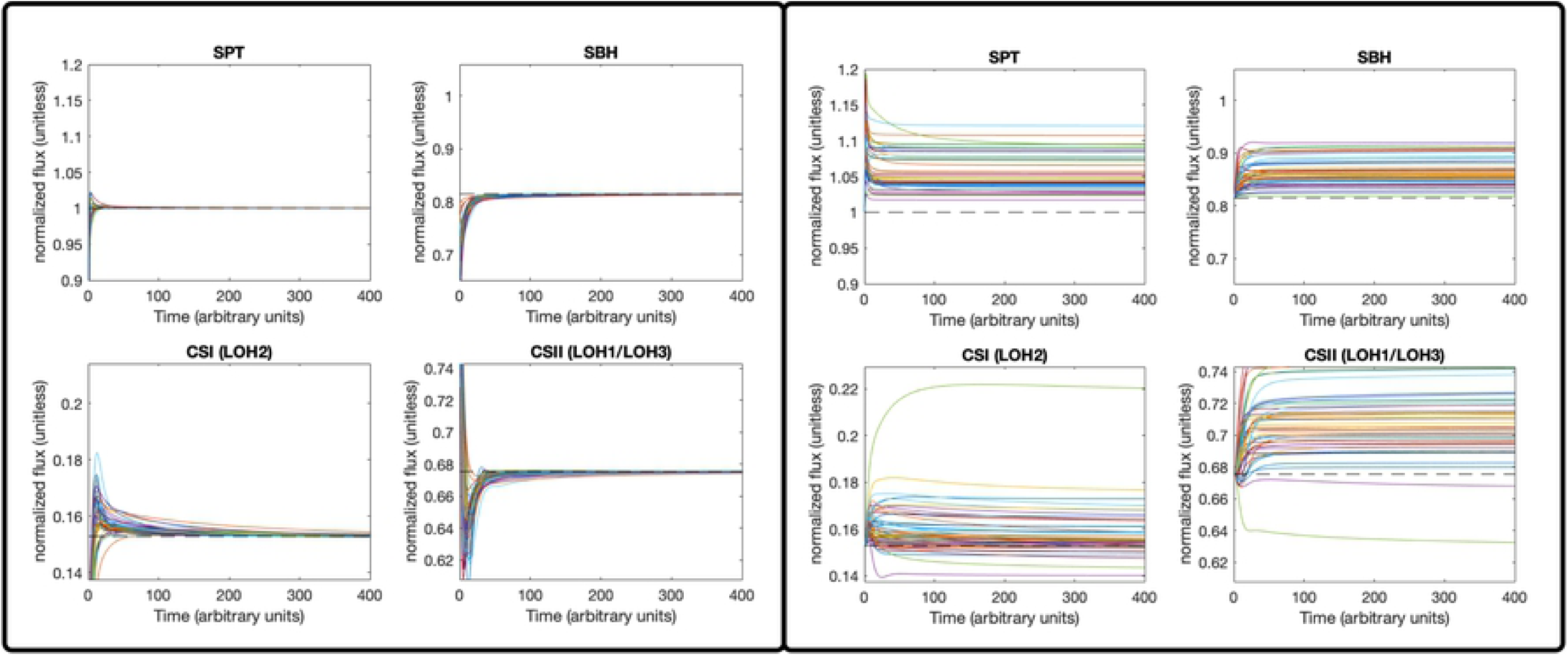
Model behavior under different conditions. (A) The flux profiles of different reactions during the reference (wild type) state. All of the models in the ensemble reach the same steady-state flux. The dashed line refers to the reference steady-state value. (B) The flux profiles of different reactions during a perturbed state. The displayed perturbation is an ORM repression. The dashed line refers to the reference steady-state value.

### The effect of perturbations on the generated model

To determine the feasibility of the metabolic network without any additional regulatory interactions, we introduced perturbations in the form of enzyme overexpression or repression and determined the predicted response of each model in the ensemble to the introduced perturbation. The predicted response was then compared with the experimentally observed response which was imposed as a ‘filtration step’. Table 2 shows the different perturbations introduced and the applied filters linked to each of them. One of the interesting observations made prior to this analysis was the differential activity in the two classes of ceramide synthase resulting from a perturbation to the level of ORMs in the system. Previous studies postulated that this differential activity potentially points to a possible regulatory interaction between ORMs and the ceramide synthases [19]. However, it was found that a large fraction of the generated models displayed this behavior without the need for any regulatory intervention, meaning that this behavior is more likely to emerge from the kinetic properties of the two enzyme classes. Furthermore, it was observed that none of the models had satisfied the filtration step requiring an increase in the concentration of the LCB sphinganine (d18:0) compared to the wild type during the overexpression of class I ceramide synthase. This observation is expected since overexpressing ceramide synthase should intuitively decrease the concentration of LCBs which are considered substrates to this enzyme. In addition, none of the models satisfied the experimental observation that the concentration of the VLCFA containing ceramide d18:1-hC24 increases during ORM overexpression. This was due to a decrease in the overall amount of LCBs produced resulting from the reduced rate through SPT. The absence of models passing these two filtration steps (see Table 1 for details) indicated that the incorporation of additional regulatory interactions was required for the model to satisfy the available experimental observations.

**Table 2.** List of perturbations and associated filtration steps used to screen the ensemble of models.

| Perturbation | Applied Filters |
| --- | --- |
| ORM RNAi* | ↑ CSI activity |
|  | ↓ CSII activity |
| ORM OE | ↓ CSI activity |
|  | ↑ CSII activity |
|  | ↓ t18:0 and d18:1-hC16 concentration* |
|  | ↑ d18:1-hC24 concentration |
| CSI OE | ↑ d18:0 concentration |
|  | ↑ “C16-cer” concentration |
|  | ↓ “C24-cer” concentration |

\*RNAi refers to inhibition of the RNA coding for ORM expression.

### Predicting the regulatory scheme of the sphingolipid pathway

To predict plausible regulatory schemes capable of reproducing all of the observed responses, we started with a set of 23 schemes that had been postulated in previous works [19,22]. Recent work in mammalian cells and yeast had found that accumulation of ceramides had an inhibitory effect on SPT activity, and that this interaction had been facilitated through ORMs [22]. In addition, previous work on the role of ORMs on the sphingolipid pathway in plants had hypothesized the possibility of a regulatory interaction between ORMs and ceramide synthases [19]. Therefore, the set of starting regulatory networks to be tested included different combinations of regulatory interactions between ORMs and the ceramide synthases and/or ceramides, as well as ceramide inhibition of SPT. These interactions were either in the form of an enzyme activation or inhibition by a component in the metabolic network. S3 File has a description of how the 23 schemes were generated and an explanation of how regulatory interactions were incorporated into the network.

The majority of the tested schemes resulted in no change to the number of filters passed compared with the starting metabolic network. The two filters discussed in the previous section had remained problematic. However, it was found that the schemes including ceramide repression of ORMs were able to satisfy all of the observed responses. It is noted that although the model captures the repression of ceramides on ORMs as a decrease in the concentration of these proteins, the observed behavior would be the same if instead ceramides repressed ORM’s binding efficiency to the regulated enzymes. Out of the 23 starting schemes, three regulatory networks all requiring ceramide repression of ORMs had passed the implemented filtration steps and were therefore considered candidates for further experiments to validate the presence of a regulatory interaction. Furthermore, it was found that the presence of an activating interaction between ORMs and CSII produced a larger number of models satisfying the observed responses. Fig 4 below displays this hypothesized regulatory network.

**Fig 4.**
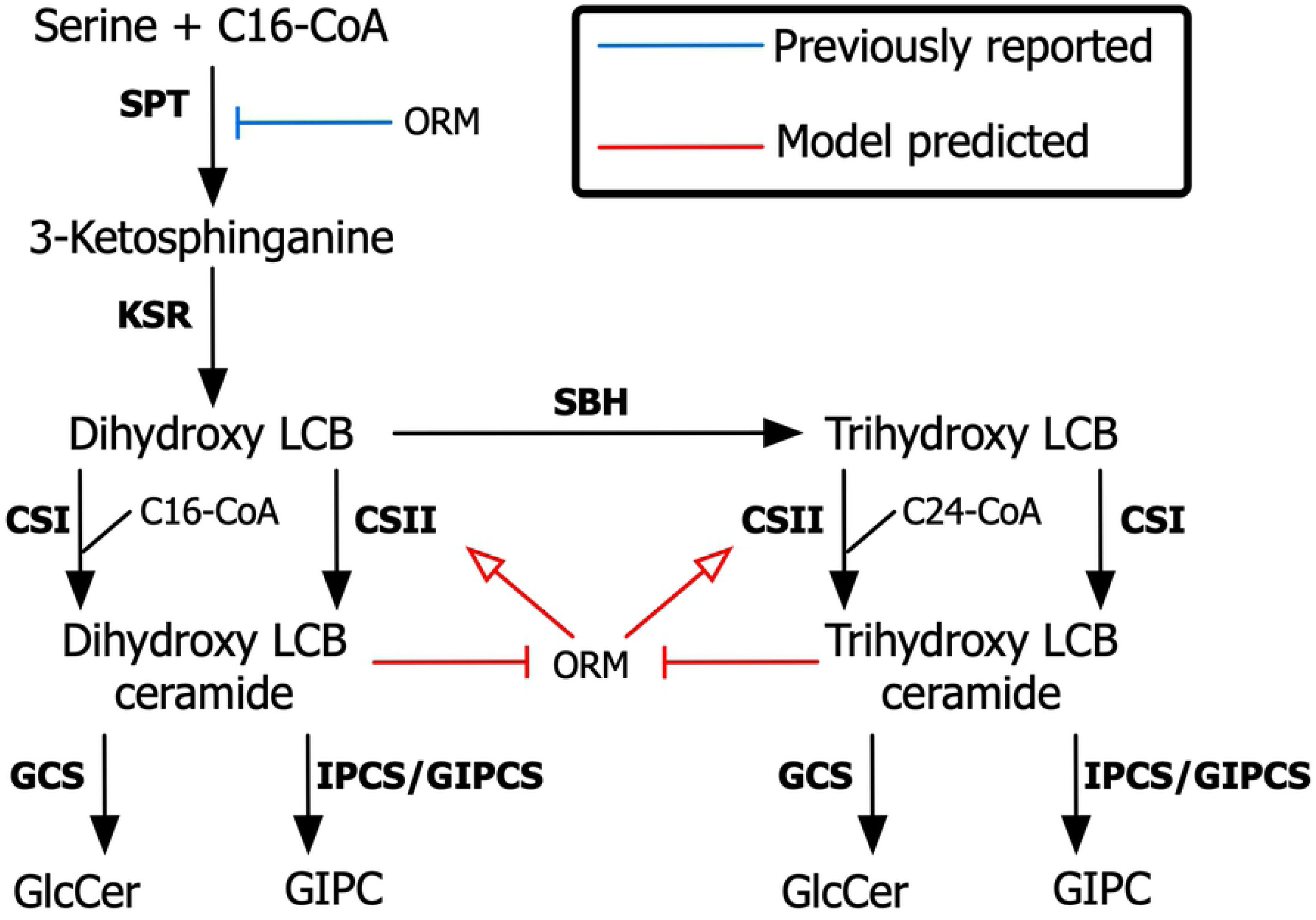
Predicted regulatory scheme of the sphingolipid pathway. A metabolic and regulatory network displaying the mechanism predicted to be responsible for the emerging observed behavior of the pathway. Ceramides repress ORM-mediated activation of CSII. Blue lines indicate regulatory interactions identified in previous studies and red lines indicate novel interactions predicted in this study.

As can be seen from Fig 4, the role of both hypothesized regulatory mechanisms is to cause an accumulation in dihydroxy-LCBs (d18:0) during CSI overexpression. The increase in ceramides resulting from the overexpression of LOH2 ceramide synthase (CSI) results in the repression of ORM-mediated activation of CSII. Analysis of the passing models showed that this does not cause a significant change in the flux through SPT. The main effect of this regulation is to dampen the activating role of ORMs on CSII. This causes the concentration of LCBs to increase, satisfying the imposed filter.

## Discussion

Sphingolipids constitute a major class of lipids in plants and play several functional and regulatory roles in plant cells [44]. The regulatory mechanisms governing the activity of the sphingolipid biosynthesis pathway are not completely understood. Specifically, even though it is well documented that ORM proteins negatively regulate SPT, questions regarding the role of these proteins in regulating different enzymes in the pathway remain unanswered [15,19,22]. In this work, we used a model guided approach to predict a regulatory scheme consistent with the pathway’s observed behavior under different conditions, the up- and downregulation of *ORM* gene expression and the overexpression of ceramide synthase class I (LOH2). The predictions made pave the way for future guided experiments and have implications in engineering crops with higher biotic stress tolerance.

Previous reports of computational analysis of sphingolipid metabolism have focused on modeling the pathway’s activity in yeast and mammalian cells under different context [30–33]. However, most of the applied frameworks either did not consider regulatory interactions or assumed a pre-defined regulatory network. A recent study of sphingolipid metabolism in yeast developed a framework to determine the regulatory effect each enzyme had on the pathway but did not incorporate the role of regulatory proteins such as ORMs [33].

In this study, a kinetic model comprised of elementary reactions describing both metabolic transformations and regulatory interactions in the sphingolipid pathway was constructed. To avoid both redundancy and overfitting, several assumptions were made to simplify the metabolic network. First, the order of reactions was assumed to be fixed based on evidence from previous works. It is possible that some of the enzymes are promiscuous in the order in which they catalyze a given reaction (e.g., LCB modification). Second, the concentration of cofactors and palmitoyl-CoA were considered to be constant both in the reference (wild type) and perturbed model. These metabolites participate in a large number of reactions in the cell and were therefore assumed to be tightly regulated (since a change in their concentration will have an effect on all the reactions they participate in). The addition of this constraint greatly reduced the number of fitting parameters required, as the cofactors are modeled to be a part of the enzyme complex. In addition, diffusion and transport rate limitations of sphingolipids entering and exiting the ER were not considered. This assumption was necessary as the transport kinetics of the pathway are not known. Finally, it was assumed that sphingolipid turnover rates were negligible compared to their rate of formation. This assumption was necessary to calculate reference steady-state fluxes. Non-stationary isotope experiments will be conducted in future studies to determine the accuracy of this assumption and the results will be used to improve the model.

The constructed network was used to generate a set of kinetic parameters that converge to the same reference steady-state fluxes. By incorporating data on how the pathway responds to the different perturbations represented in Table 2, different regulatory schemes were tested to determine which ones were consistent with the observed data. The experimentally observed pathway responses to different perturbations [enzyme over-expression (OE)/downregulation] were incorporated into the framework as filtration steps that the generated models needed to satisfy.

This was implemented using a modified ensemble modeling framework (see Materials and Methods). Namely, the process of model screening was automated through the addition of a new module which filters through the initial ensemble of kinetic parameters. Furthermore, the incorporation of an activating regulatory mechanism was amended to ensure the generation of models consistent with the reference steady-state.

Starting with only the known regulatory interaction between ORMs and SPT, it was found that this mechanism alone was sufficient to obtain the observed differential activity of the two classes of ceramide synthases during ORM OE or repression. This observation was counterintuitive as both enzymes use LCBs as their substrate. The model showed that this behavior could be explained solely based on the kinetics of the two classes of enzymes and does not require any further regulation of either ceramide synthase. However, this scheme was still incomplete, as it did not satisfy other observed behaviors in the pathway. Namely, the observation that LCBs accumulate during *LOH2* (CSI) OE required additional regulatory mechanisms to be incorporated into the network. By testing a set of plausible regulatory schemes, it was observed that the addition of a ceramide-ORM inhibitory interaction and an ORM activation of class II ceramide synthases was required for all observed filtration steps to be satisfied. This scheme is depicted in Fig 4. In this scenario, the prediction of the model indicates two layers of regulation under normal conditions, (1) ORMs activate the synthesis of C24-containing ceramides by LOH1/LOH3 (CSII) and (2) the accumulation of ceramides represses this ORM-mediated activation on CSII. The first prediction is consistent with a previous study where overexpression of ORMs resulted in increased activity of class II CS that preferentially use VLCFA and trihydroxy LCBs as substrates [19]. However, the effect of the accumulation of ceramides in this regulation has not been reported. This repression by ceramides could also be explained by a competitive inhibition of ceramide synthase.

In mammalian cells, it has been shown that ceramides, or downstream metabolites, are involved in regulating the expression of ORM proteins, ceramide accumulation leads to increased ORM protein levels [45]. In addition, ceramides mediate SPT inhibition by ORMs [22]. The role of ORMDL proteins in the regulation of ceramide synthesis was tested using mammalian cell cultures. However, in this system ORMDLs do not regulate ceramide synthases [20]. The biological implications of the ceramide-ORM-CSII regulation depicted in the present study could be specific to plants where ceramides containing VLCFA and trihydroxy LCBs are very abundant compared to mammalian cells. This regulation could be relevant under bacterial and fungal infections where differential accumulation of C16-containing ceramides and VLCFA-containing ceramides might be related to the defense response [6]. Therefore, future experimental work will focus on verifying the interaction of ORMs-LOHs and the ORM-mediated activation of CSII. Moreover, further studies will be conducted using the ceramide synthase inhibitor fumonisin B1 (FB1). This fungal toxin acts preferentially on LOH1 [46] promoting the accumulation of C16 ceramides though LOH2. These results will be compared with the LOH2 OE scheme presented in this work.

The main strength of using the ensemble modeling framework to approach such challenges is its ability to eliminate regulatory schemes which appear to be plausible on first sight. As demonstrated in this work, the analysis significantly reduced the number of plausible regulatory schemes to be tested experimentally. Furthermore, due to the computational intractability of testing all combinations of regulatory interactions in the pathway, it is possible that other regulatory networks can also satisfy all filtration steps applied in this study. For example, ORMs (or other components) could also be involved in regulating other downstream enzymes like GCS. As more experiments are carried out on different Arabidopsis mutants and transgenic lines, the number of filtration steps will increase, potentially leading to fewer schemes that satisfy all observations. Furthermore, experiments aimed at looking for regulatory interactions between ORMs and different parts of the network might unravel additional interactions that were not tested in this work. These schemes can easily be incorporated into the model as the metabolic reactions have already been constructed. This highlights the need for future experimental work to test the predictions of this study, which will allow the model to be updated in a more systematic manner. We emphasize the importance of such model-driven approaches in addressing such problems as the one in this study, where the aim of the predictions is (i) to reduce the solution space that the experimentalist has to cover and (ii) to pin-point parts of the pathway where more data is needed to obtain a more accurate model. In this regard, this work can be considered as the first step in a continuously updated design-test-refine cycle aimed at obtaining an accurate mechanistic understanding of the behavior of the sphingolipid pathway in plants.

## Methods

### Sphingolipid compositions

The concentrations of LCB, ceramides, GlcCer, and GIPC in 12 to15-day-old wild-type (Columbia-0) seedlings were obtained from a previous study [16].

### Growth rate

Arabidopsis seedlings were grown on Murashige and Skoog (MS) medium supplemented with 1% sucrose and 0.8% agar (pH 5.7) with 16 h light (100μmol/ m^-2^ s^-1^) 8 h dark conditions at 22°C. Three replicates, consisting of ten seedlings, were sampled at each time point (5 d, 10 d, 15d and 20 d). The seedlings were freeze-dried, and the dry weight recorded to determine the growth rate.

### Ensemble Modeling

EM describes both the metabolic and regulatory state of a pathway as a set of elementary steps obeying mass-action kinetics. This allows the construction of multiple sets of regulatory schemes, which can be subsequently screened to determine the thermodynamic and kinetic feasibility of each network. This section contains a brief description of how the set of kinetic parameters can be obtained for a unimolecular enzymatic reaction with no regulation. Complex reaction schemes follow the same general procedure but contain more elementary steps.

#### Determining steady-state flux distributions

Parsimonious Flux Balance Analysis (pFBA) [47] was used to generate the steady state flux distribution. pFBA is analogous to FBA but adds an outer objective that minimizes the sum of all reaction fluxes. Objective tilting [48] was used to formulate both objectives in one function as shown below.

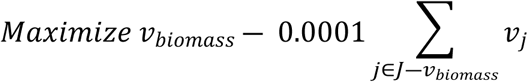

*subject to*

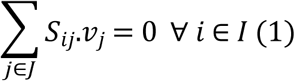

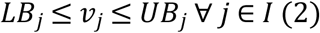

Where *I* and *J* are the sets of metabolites and reactions in the model, respectively. *S*_*ij*_ is the stoichiometric coefficient of metabolite *i* in reaction *j* and *v*_*j*_ is the flux value of reaction *j*. Parameters *LB*_*j*_ and *UB*_*j*_ denote the minimum and maximum allowable fluxes for reaction *j*, respectively. *v*_*biomass*_ is the flux of the biomass reaction which mimics the cellular growth rate.

The cellular composition of each of the sphingolipid components was experimentally measured and used to establish their stoichiometric coefficients of the biomass equation. Since the majority of ceramides and complex sphingolipids contained either 16 or 24 carbon fatty acids, other acyl-CoA lengths were pooled together with one of the two groups depending on whether they contained more or less than twenty carbons. The growth rate was also determined experimentally to constrain the rate of the biomass reaction. Subsequently, by assuming negligible sphingolipid turnover rates, pFBA was used to calculate the reaction rates for each enzymatic reaction in the network.

#### Generating initial ensemble of models

The elementary steps for a unimolecular enzymatic reaction can be written as:

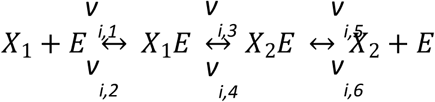

where, enzyme E_i_ converts metabolite X_i_ into X_i+1_. Each step is reversible and has a rate of v_i,2j-1_ in the forward direction and v_i,2j_ in the reverse direction. Since each elementary step follows mass-action kinetics, the reaction rate can be expressed as the product of the reactant concentrations and a rate constant. Therefore, the rate of the forward reaction of the first step can be written as:

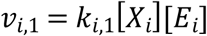

where, k_i,1_ is the rate constant associated with the forward reaction of the first step. This equation is subsequently normalized by the reference steady-state metabolite concentration and the total enzyme concentration to yield:

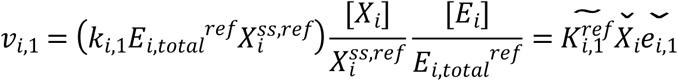

where, E^ref^_i,total_ is the total enzyme concentration and X_i_^ss,ref^ is the steady state metabolite concentration, both at the reference steady state. This equation is subsequently converted to the log-linear form:

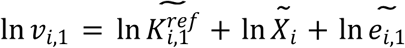

During steady-state, X_i_ becomes 1, therefore the ln(X_i_) term can be omitted. This yields:

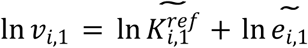

Next, the reversibility of each reaction step (R_i,j_) is sampled in order to determine the rate of each elementary reaction v_i,j_

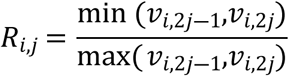

R_i,j_ can take on any value between 0 and 1 (where 0 is an irreversible reaction step) and is further constrained by the gibbs free energy of the reaction as follows

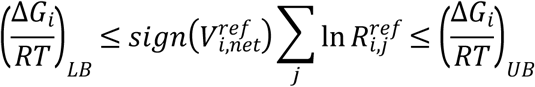

The reaction rates for each step (v_i,j_) can then be determined from the net enzymatic reaction rate (V^ref^_i,net_)

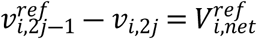

Finally, the enzyme fractions for each elementary step (e_i,j_) are sampled between 0 and 1 in order to calculate the kinetic parameter (K _i,j_ ^ref^). An additional constraint is imposed to ensure that the sum of enzyme fractions for each reaction is unity.

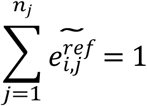

This procedure is repeated thousands of times to generate an ensemble of kinetic parameters that all reach the predefined reference steady-state. Next, a system of ordinary differential equations describing the metabolic network is solved to obtain the net steady-state fluxes.

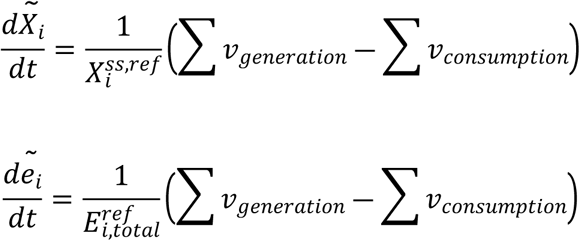

A modified version of the MATLAB implementation developed by Tran et. al. [26] was used to carry out the simulations in this work.

#### Model screening

After the initial ensemble of models are generated, a number of enzymatic perturbations are introduced to analyze each model’s response. The perturbations take on the form of enzyme overexpression, inhibition, or knockout and are formulated as follows

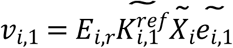

Where E_i,r_ represents the introduced fold change in enzyme expression compared to the reference state. Although the original MATLAB implementation [26] incorporated both inhibition and activation regulatory mechanisms, it was observed that no models were generated when activation was incorporated into the regulatory network. Therefore, modifications were implemented to ensure that the mass balances of the introduced activator-enzyme complexes are satisfied. Furthermore, the fluxes of the activated pathway(s) were integrated with the flux going through the main reaction to ensure an accurate response to changes in the concentration of the activator.

After each perturbation, the predicted enzymatic response is compared with the experimentally observed behavior. Models resulting in contradictory enzymatic behavior are disregarded. In this manner, the initial set of models is filtered through multiple rounds of perturbations, until only a small set of highly predictive models remain. A new module was added to the original implementation to automate the screening process. Perturbations with experimentally known responses were incorporated into the module in order to automatically screen the initial ensemble of models. The output is a subset of models passing all filtration steps. It is noted that the order in which the screenings are introduced does not affect the final outcome.

#### Constructing multiple sets of possible regulatory networks

As described earlier, regulatory events can be described as elementary reaction steps [49]. The regulatory network is input as a stoichiometric matrix (Sreg) similar in size to the S matrix used to describe the metabolic network. The element Sreg_i,j_ describes the type of regulation metabolite i confers on enzyme j, and can either be activating or inhibitory. Therefore, by constructing multiple Sreg matrices corresponding to different possible regulatory schemes, the EM approach can be applied on each regulatory network individually. Furthermore, after generating and filtering the set of kinetic parameters for each network, the regulatory scheme passing all the screening steps is hypothesized to be the most accurately representative of the biological system.

## Acknowledgement

Funding to support this work was provided by a National Science Foundation grant MCB 1818297 to EBC and RS. We thank Dr. Jonathan E. Markham, Dr. Gongshe Han, and Dr. Teresa M. Dunn for their valuable input.

## Author contributions

AA, AGS, EBC, and EBC conceived the idea of the project. A.A. reconstructed the model and conducted computational analysis. AGS conducted growth and sphingolipid profile measuremetns. All authors designed research, wrote and edited the manuscript.

## Supporting information

**S1 File. Sphingolipid biosynthesis metabolic network**.

**S2 File. Standard Gibbs free energy of reactions in the network**.

**S3 File. Set of regulatory schemes tested in this work**.

## References

1. Luttgeharm KD, Kimberlin AN, Cahoon EB. Plant Sphingolipid Metabolism and Function BT - Lipids in Plant and Algae Development. In: Nakamura Y, Li-Beisson Y, editors. Cham: Springer International Publishing; 2016. p. 249–86. Available from: https://doi.org/10.1007/978-3-319-25979-6_11

2. Cacas J-L, Buré C, Grosjean K, Gerbeau-Pissot P, Lherminier J, Rombouts Y, et al. Revisiting Plant Plasma Membrane Lipids in Tobacco: A Focus on Sphingolipids. Plant Physiol [Internet]. 2015/10/30. 2016 Jan;170(1):367–84. Available from: https://pubmed.ncbi.nlm.nih.gov/26518342

3. Michaelson L V, Napier JA, Molino D, Faure J-D. Plant sphingolipids: Their importance in cellular organization and adaption. Biochim Biophys Acta [Internet]. 2016/04/13. 2016 Sep;1861(9 Pt B):1329–35. Available from: https://pubmed.ncbi.nlm.nih.gov/27086144

4. Melser S, Batailler B, Peypelut M, Poujol C, Bellec Y, Wattelet-Boyer V, et al. Glucosylceramide Biosynthesis is Involved in Golgi Morphology and Protein Secretion in Plant Cells. Traffic [Internet]. 2010 Apr 1;11(4):479–90. Available from: https://doi.org/10.1111/j.1600-0854.2009.01030.x

5. Liang H, Yao N, Song JT, Luo S, Lu H, Greenberg JT. Ceramides modulate programmed cell death in plants. Genes Dev [Internet]. 2003/10/16. 2003 Nov 1;17(21):2636–41. Available from: https://pubmed.ncbi.nlm.nih.gov/14563678

6. Magnin-Robert M, Le Bourse D, Markham J, Dorey S, Clément C, Baillieul F, et al. Modifications of Sphingolipid Content Affect Tolerance to Hemibiotrophic and Necrotrophic Pathogens by Modulating Plant Defense Responses in Arabidopsis. Plant Physiol [Internet]. 2015/09/16. 2015 Nov;169(3):2255–74. Available from: https://pubmed.ncbi.nlm.nih.gov/26378098

7. Hanada K. Serine palmitoyltransferase, a key enzyme of sphingolipid metabolism. Biochim Biophys Acta - Mol Cell Biol Lipids [Internet]. 2003;1632(1):16–30. Available from: http://www.sciencedirect.com/science/article/pii/S1388198103000593

8. Markham JE, Molino D, Gissot L, Bellec Y, Hématy K, Marion J, et al. Sphingolipids Containing Very-Long-Chain Fatty Acids Define a Secretory Pathway for Specific Polar Plasma Membrane Protein Targeting in *Arabidopsis*; Plant Cell [Internet]. 2011 Jun 1;23(6):2362 LP – 2378. Available from: http://www.plantcell.org/content/23/6/2362.abstract

9. Ternes P, Feussner K, Werner S, Lerche J, Iven T, Heilmann I, et al. Disruption of the ceramide synthase LOH1 causes spontaneous cell death in Arabidopsis thaliana. New Phytol [Internet]. 2011 Dec 1;192(4):841–54. Available from: https://doi.org/10.1111/j.1469-8137.2011.03852.x

10. Luttgeharm KD, Chen M, Mehra A, Cahoon RE, Markham JE, Cahoon EB. Overexpression of Arabidopsis Ceramide Synthases Differentially Affects Growth, Sphingolipid Metabolism, Programmed Cell Death, and Mycotoxin Resistance. Plant Physiol [Internet]. 2015/08/14. 2015 Oct;169(2):1108–17. Available from: https://pubmed.ncbi.nlm.nih.gov/26276842

11. Msanne J, Chen M, Luttgeharm KD, Bradley AM, Mays ES, Paper JM, et al. Glucosylceramides are critical for cell-type differentiation and organogenesis, but not for cell viability in Arabidopsis. Plant J [Internet]. 2015 Oct;84(1):188–201. Available from: https://pubmed.ncbi.nlm.nih.gov/26313010

12. Mortimer JC, Scheller HV. Synthesis and Function of Complex Sphingolipid Glycosylation. Trends Plant Sci [Internet]. 2020 Jun 1;25(6):522–4. Available from: https://doi.org/10.1016/j.tplants.2020.03.007

13. Kimberlin AN, Majumder S, Han G, Chen M, Cahoon RE, Stone JM, et al. Arabidopsis 56-amino acid serine palmitoyltransferase-interacting proteins stimulate sphingolipid synthesis, are essential, and affect mycotoxin sensitivity. Plant Cell [Internet]. 2013/11/08. 2013 Nov;25(11):4627–39. Available from: https://pubmed.ncbi.nlm.nih.gov/24214397

14. Breslow DK, Collins SR, Bodenmiller B, Aebersold R, Simons K, Shevchenko A, et al. Orm family proteins mediate sphingolipid homeostasis. Nature [Internet]. 2010 Feb 25;463(7284):1048–53. Available from: https://pubmed.ncbi.nlm.nih.gov/20182505

15. Li J, Yin J, Rong C, Li K-E, Wu J-X, Huang L-Q, et al. Orosomucoid Proteins Interact with the Small Subunit of Serine Palmitoyltransferase and Contribute to Sphingolipid Homeostasis and Stress Responses in Arabidopsis. Plant Cell [Internet]. 2016/12/06. 2016 Dec;28(12):3038–51. Available from: https://pubmed.ncbi.nlm.nih.gov/27923879

16. Gonzalez Solis A, Han G, Gan L, Liu Y, Markham JE, Cahoon RE, et al. Unregulated Sphingolipid Biosynthesis in Gene-Edited Arabidopsis ORM Mutants Results in Nonviable Seeds with Strongly Reduced Oil Content. Plant Cell [Internet]. 2020 Jan 1;tpc.00015.2020. Available from: http://www.plantcell.org/content/early/2020/06/11/tpc.20.00015.abstract

17. Clarke BA, Majumder S, Zhu H, Lee YT, Kono M, Li C, et al. The Ormdl genes regulate the sphingolipid synthesis pathway to ensure proper myelination and neurologic function in mice. Elife [Internet]. 2019 Dec 27;8:e51067. Available from: https://pubmed.ncbi.nlm.nih.gov/31880535

18. Han G, Gupta SD, Gable K, Bacikova D, Sengupta N, Somashekarappa N, et al. The ORMs interact with transmembrane domain 1 of Lcb1 and regulate serine palmitoyltransferase oligomerization, activity and localization. Biochim Biophys Acta - Mol Cell Biol Lipids [Internet]. 2019;1864(3):245–59. Available from: http://www.sciencedirect.com/science/article/pii/S138819811830194X

19. Kimberlin AN, Han G, Luttgeharm KD, Chen M, Cahoon RE, Stone JM, et al. ORM Expression Alters Sphingolipid Homeostasis and Differentially Affects Ceramide Synthase Activity. Plant Physiol [Internet]. 2016/08/09. 2016 Oct;172(2):889–900. Available from: https://pubmed.ncbi.nlm.nih.gov/27506241

20. Siow DL, Wattenberg BW. Mammalian ORMDL proteins mediate the feedback response in ceramide biosynthesis. J Biol Chem [Internet]. 2012/10/12. 2012 Nov 23;287(48):40198–204. Available from: https://pubmed.ncbi.nlm.nih.gov/23066021

21. Siow D, Sunkara M, Dunn TM, Morris AJ, Wattenberg B. ORMDL/serine palmitoyltransferase stoichiometry determines effects of ORMDL3 expression on sphingolipid biosynthesis. J Lipid Res [Internet]. 2015/02/17. 2015 Apr;56(4):898–908. Available from: https://pubmed.ncbi.nlm.nih.gov/25691431

22. Davis DL, Gable K, Suemitsu J, Dunn TM, Wattenberg BW. The ORMDL/Orm-serine palmitoyltransferase (SPT) complex is directly regulated by ceramide: Reconstitution of SPT regulation in isolated membranes. J Biol Chem [Internet]. 2019/01/30. 2019 Mar 29;294(13):5146–56. Available from: https://pubmed.ncbi.nlm.nih.gov/30700557

23. Islam MM, Saha R. Computational Approaches on Stoichiometric and Kinetic Modeling for Efficient Strain Design BT - Synthetic Metabolic Pathways: Methods and Protocols. In: Jensen MK, Keasling JD, editors. New York, NY: Springer New York; 2018. p. 63–82. Available from: https://doi.org/10.1007/978-1-4939-7295-1_5

24. Zielinski DC, Palsson BØ. Kinetic Modeling of Metabolic Networks BT - Systems Metabolic Engineering. In: Wittmann C, Lee SY, editors. Dordrecht: Springer Netherlands; 2012. p. 25–55. Available from: https://doi.org/10.1007/978-94-007-4534-6_2

25. Rohwer JM. Kinetic modelling of plant metabolic pathways. J Exp Bot [Internet]. 2012 Mar 1;63(6):2275–92. Available from: https://doi.org/10.1093/jxb/ers080

26. Tran LM, Rizk ML, Liao JC. Ensemble modeling of metabolic networks. Biophys J. 2008;95(12):5606–17.

27. Rizk ML, Liao JC. Ensemble modeling and related mathematical modeling of metabolic networks. J Taiwan Inst Chem Eng. 2009;40(6):595–601.

28. Rizk ML, Liao JC. Ensemble modeling for aromatic production in Escherichia coli. PLoS One [Internet]. 2009 Sep 4;4(9):e6903–e6903. Available from: https://pubmed.ncbi.nlm.nih.gov/19730732

29. Khazaei T, McGuigan A, Mahadevan R. Ensemble modeling of cancer metabolism. Front Physiol [Internet]. 2012 May 16;3:135. Available from: https://pubmed.ncbi.nlm.nih.gov/22623918

30. Gupta S, Maurya MR, Merrill Jr AH, Glass CK, Subramaniam S. Integration of lipidomics and transcriptomics data towards a systems biology model of sphingolipid metabolism. BMC Syst Biol [Internet]. 2011;5(1):26. Available from: https://doi.org/10.1186/1752-0509-5-26

31. Alvarez-Vasquez F, Sims KJ, Cowart LA, Okamoto Y, Voit EO, Hannun YA. Simulation and validation of modelled sphingolipid metabolism in Saccharomyces cerevisiae. Nature [Internet]. 2005;433(7024):425–30. Available from: https://doi.org/10.1038/nature03232

32. Wronowska W, Charzynska A, Nienaltowski K, Gambin A. Computational modeling of sphingolipid metabolism. BMC Syst Biol [Internet]. 2015;9(1):47. Available from: https://doi.org/10.1186/s12918-015-0176-9

33. Savoglidis G, Santos A, Riezman I, Angelino P, Riezman H, Hatzimanikatis V. A method for analysis and design of metabolism using metabolomics data and kinetic models: Application on lipidomics using a novel kinetic model of sphingolipid metabolism. Metab Eng. 2016 Apr 22;37.

34. Michaelson L V, Zäuner S, Markham JE, Haslam RP, Desikan R, Mugford S, et al. Functional characterization of a higher plant sphingolipid Delta4-desaturase: defining the role of sphingosine and sphingosine-1-phosphate in Arabidopsis. Plant Physiol [Internet]. 2008/10/31. 2009 Jan;149(1):487–98. Available from: https://pubmed.ncbi.nlm.nih.gov/18978071

35. Markham J, Li J, Cahoon E, Jaworski J. Separation and Identification of Major Plant Sphingolipid Classes from Leaves. J Biol Chem. 2006 Sep 1;281:22684–94.

36. Chen M, Markham JE, Dietrich CR, Jaworski JG, Cahoon EB. Sphingolipid long-chain base hydroxylation is important for growth and regulation of sphingolipid content and composition in Arabidopsis. Plant Cell [Internet]. 2008/07/08. 2008 Jul;20(7):1862–78. Available from: https://pubmed.ncbi.nlm.nih.gov/18612100

37. König S, Feussner K, Schwarz M, Kaever A, Iven T, Landesfeind M, et al. Arabidopsis mutants of sphingolipid fatty acid α-hydroxylases accumulate ceramides and salicylates. New Phytol [Internet]. 2012 Dec 1;196(4):1086–97. Available from: https://doi.org/10.1111/j.1469-8137.2012.04351.x

38. Habel A, Sperling P, Bartram S, Heinz E, Boland W. Conformational Studies on the Δ8(E,Z)-Sphingolipid Desaturase from Helianthus annuus with Chiral Fluoropalmitic Acids As Mechanistic Probes. J Org Chem [Internet]. 2010 Aug 6;75(15):4975–82. Available from: https://doi.org/10.1021/jo100542q

39. Chen M, Markham JE, Cahoon EB. Sphingolipid Δ8 unsaturation is important for glucosylceramide biosynthesis and low-temperature performance in Arabidopsis. Plant J [Internet]. 2012 Mar 1;69(5):769–81. Available from: https://doi.org/10.1111/j.1365-313X.2011.04829.x

40. Rennie EA, Ebert B, Miles GP, Cahoon RE, Christiansen KM, Stonebloom S, et al. Identification of a Sphingolipid α-Glucuronosyltransferase That Is Essential for Pollen Function in *Arabidopsis*; Plant Cell [Internet]. 2014 Aug 1;26(8):3314 LP – 3325. Available from: http://www.plantcell.org/content/26/8/3314.abstract

41. Orth JD, Thiele I, Palsson BØ. What is flux balance analysis? Nat Biotechnol [Internet]. 2010 Mar;28(3):245–8. Available from: https://pubmed.ncbi.nlm.nih.gov/20212490

42. Noor E, Haraldsdóttir HS, Milo R, Fleming RMT. Consistent estimation of Gibbs energy using component contributions. PLoS Comput Biol [Internet]. 2013/07/11. 2013;9(7):e1003098–e1003098. Available from: https://pubmed.ncbi.nlm.nih.gov/23874165

43. Noor E, Flamholz A, Liebermeister W, Bar-Even A, Milo R. A note on the kinetics of enzyme action: A decomposition that highlights thermodynamic effects. FEBS Lett [Internet]. 2013 Sep 2;587(17):2772–7. Available from: https://doi.org/10.1016/j.febslet.2013.07.028

44. Chen M, Cahoon EB, Saucedo-García M, Plasencia J, Gavilanes-Ruíz M. Plant Sphingolipids: Structure, Synthesis and Function BT - Lipids in Photosynthesis: Essential and Regulatory Functions. In: Wada H, Murata N, editors. Dordrecht: Springer Netherlands; 2009. p. 77–115. Available from: https://doi.org/10.1007/978-90-481-2863-1_5

45. Gupta SD, Gable K, Alexaki A, Chandris P, Proia RL, Dunn TM, et al. Expression of the ORMDLS, modulators of serine palmitoyltransferase, is regulated by sphingolipids in mammalian cells. J Biol Chem [Internet]. 2014/11/13. 2015 Jan 2;290(1):90–8. Available from: https://pubmed.ncbi.nlm.nih.gov/25395622

46. Luttgeharm KD, Cahoon EB, Markham JE. Substrate specificity, kinetic properties and inhibition by fumonisin B1 of ceramide synthase isoforms from Arabidopsis. Biochem J [Internet]. 2016 Feb 24;473(5):593–603. Available from: https://doi.org/10.1042/BJ20150824

47. Lewis NE, Hixson KK, Conrad TM, Lerman JA, Charusanti P, Polpitiya AD, et al. Omic data from evolved E. coli are consistent with computed optimal growth from genome-scale models. Mol Syst Biol [Internet]. 2010 Jan 1;6(1):390. Available from: https://doi.org/10.1038/msb.2010.47

48. Feist AM, Zielinski DC, Orth JD, Schellenberger J, Herrgard MJ, Palsson BØ. Model-driven evaluation of the production potential for growth-coupled products of Escherichia coli. Metab Eng [Internet]. 2010;12(3):173–86. Available from: http://www.sciencedirect.com/science/article/pii/S1096717609000895

49. Khodayari A, Maranas CD. A genome-scale Escherichia coli kinetic metabolic model k-ecoli457 satisfying flux data for multiple mutant strains. Nat Commun. 2016;7.

